# Sedimentary ancient DNA metagenomic analysis provides new insights into farming in central Norway from the Bronze Age to late Medieval period

**DOI:** 10.1101/2025.06.19.660509

**Authors:** Renato La Torre, Paula Kosel, Vanessa C. Bieker, Kristine Bakke Westergaard, Jonas Lien Mykkeltvedt, Kathinka Louisa Yildirim, Martin Seiler, Bente Philippsen, Richard I Macphail, Rebecca J S Cannell, Lene Synnøve Halvorsen, Astrid Brønseth Lorentzen, Raymond Sauvage, Michael D. Martin, Heidi Mjelva Breivik, Sarah L. F. Martin

## Abstract

Sedimentary ancient DNA (sedaDNA) has been proposed as a key methodology for reconstructing paleoclimates and biodiversity over time. To a lesser extent, it has been explored as a complementary tool for reconstructing human-driven local environmental changes over time, such as those explored in open air archaeological sites. Our study employs a sedaDNA metagenomic approach to investigate land use and environmental change at the archaeological site of Torgårdsletta in central Norway, spanning from the Bronze Age through the Medieval period. Stratigraphic sediment samples reveal temporal shifts in plant, animal and microbial communities, reflecting evolving human practices and climatic conditions. Progression through the layers indicate signs or land clearance, cultivation and animal husbandry in the region. Notably, a significant reduction in microbial and plant diversity during periods of climatic upheaval—such as the AD 536–540 volcanic event—correlates with landscape and societal adjustments. The findings demonstrate that sedaDNA complements traditional proxies, providing high-resolution insights into past land use, environmental interactions, and societal organization. The successful extraction of ancient genetic material underscores sedaDNA’s potential to reconstruct dynamic prehistoric landscapes and anthropogenic impacts, offering further potential for understanding long-term human-environment relationships in Scandinavia.

## 1. INTRODUCTION

Sedimentary ancient DNA (sedaDNA), obtained directly from sediment samples, has experienced a surge in interest over the last decade. Its analysis has proven capable of detecting taxa in samples up to two million years old^1^, showcasing remarkable potential for reconstructing past ecosystem changes across diverse organisms, from plants and animals to fungi and microorganisms. While the majority of sedaDNA studies have focused on ecological questions^1–7^, its application in archaeological contexts is gaining momentum, with researchers exploring its potential for routine use in the field^8,9^.

Although early archaeological sedaDNA studies often focused on sediments from caves and rockshelters^8,9^, successful applications at open-air sites are also emerging^10,11^. However, obtaining sedaDNA of sufficient quality for analysis from archaeological sites poses significant challenges^12^. The persistence of these biomolecules hinges on favorable preservation conditions, as heat, oxygen, and water promote DNA degradation. Given the geographic and temporal variability in environmental conditions, sedaDNA quality will inevitably differ across archaeological sites. As such, it is important to continue applying sedaDNA methods in numerous regions and site types to increase our understanding of preservation condition expectations and evaluate the implementation of sedaDNA in routine archaeological excavations along with other currently used proxies.

SedaDNA analyses offer a valuable complement to traditional scientific methods utilized in archaeology, such as stratigraphic observations and pollen and macrofossil analysis, by providing information on taxa that are poorly preserved in the sedimentary record. By integrating genetic data into archaeological excavations, researchers can achieve a more nuanced understanding of past human behavior and environmental interactions.

Norway, with its rich archaeological heritage spanning nearly 12,000 years, presents a unique challenge due to climatic and geological conditions that often lead to poor preservation of organic materials. The application of sedaDNA analysis in Norway could unlock new discoveries and interpretations, shaping future research by enhancing our understanding of the complex relationships between ancient human populations and their environment, including animals and food resources.

This study focuses on Torgårdsletta, a late glacial moraine formation located 14 km outside Trondheim in central Norway (Figure 1). Modern archaeological rescue excavations at this site, prompted by the planned transition from an agricultural to an industrial landscape, have revealed extensive evidence of settlement, farming, and graves dating from the Bronze Age (c. 1000 BC) through the medieval period (1050-1550 AD)^13,14^. Pre-excavation surveys identified features indicative of farming activity (e.g. houses and cooking pits) contemporary with other sites along the ridge, and the stratigraphy revealed layers of buried agricultural soils and peat horizons. Preservation conditions at the site resulted in no bone fragments being found, thus to date it has only been hypothesized which domesticated animals were utilized. The distinct stratification, rich sediment build-up, and character of these layers prompted us to investigate the dynamics of agricultural activity through time in greater detail. To address the changes in land use over time, and to evaluate the potential of sedaDNA as a routine proxy in future rescue archaeology projects, we applied a metagenomic approach to sedaDNA recovered from the Torgårdsletta site. Our study aims to assess the feasibility of obtaining sedaDNA from the soil of a typical prehistoric farm site, representing an open-air context, while taxonomically characterizing plant, animal, and microbial diversity at the site through time to understand the dynamics of humans and their surrounding environment. In addition to the sedaDNA, we integrate findings from radiocarbon dating, soil micromorphology, pollen and macrofossil analysis.

**Figure 1.**
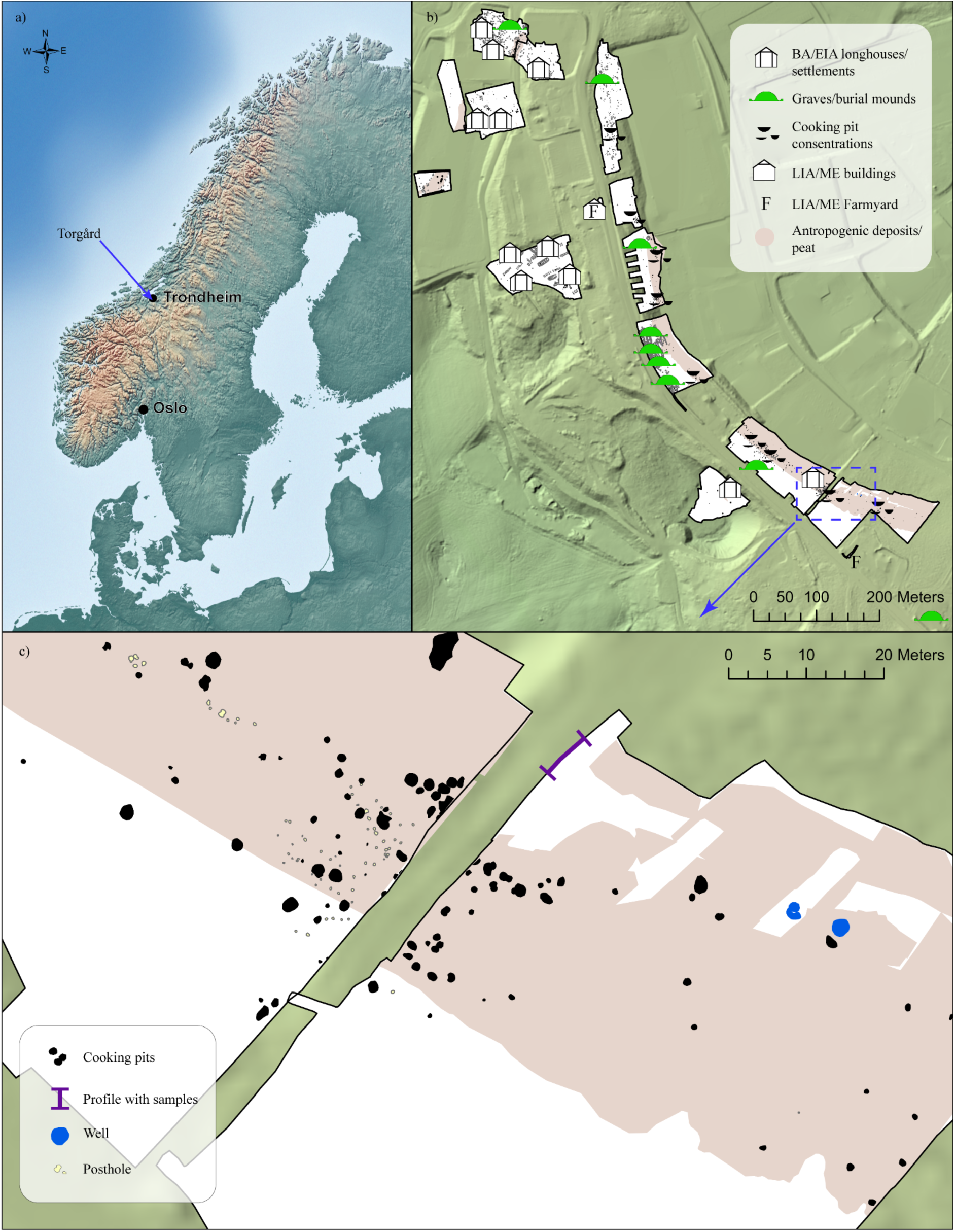
A) Map showing location of Torgård archaeological site; B) Overview of key archaeological finds at Torgårdsletta; C) Precise location of the profile examined in this study.

## 2. MATERIALS AND METHODS

### 2.1. The site: Torgård Østre

Archaeological and topographical observations at Torgårdsletta (Figure 1) recorded in the 18th and 19th centuries indicate that the entire ridge held a large number of burial mounds that indicate considerable prehistoric activity. Several finds are recorded from the area, which predominantly dates the activity to the Nordic Iron Age (500 BC-AD 1050). The most famous is a Vendel-type 7th-century elite equestrian warrior grave (discovered in 1933), which suggests that in the early Late Iron Age (AD 550 – 1050), the region was controlled by a person involved in over-regional Northern European elite networks^15^. The specific area of interest to this study is the eastern slope of the moraine ridge, Torgård Østre, which was part of a large archaeological rescue excavation performed in 2023 by the NTNU University Museum. The features recognized in the pre-excavation surveys, indicated farming activity contemporary with other sites along the ridge. The stratigraphy included layers of buried agricultural soils and peat horizons. Among the finds from the site were three Early Medieval (1050-1150 AD) wooden wells, the remains of wooden fence works, and cooking pits (see Figure 1C).

Radiocarbon dates (Figure 2) suggest that the lower layer was first formed c. BC 1411 – 1279 in the Bronze Age. The dates from the upper portion of the topmost peat layer dates to c. 1030 – 1155 AD in the Early Medieval period. Including another agricultural layer accumulating on top of the peat, this indicates at least 3,000 years of soil formation and intensive cultivation in several phases.

**Figure 2.**
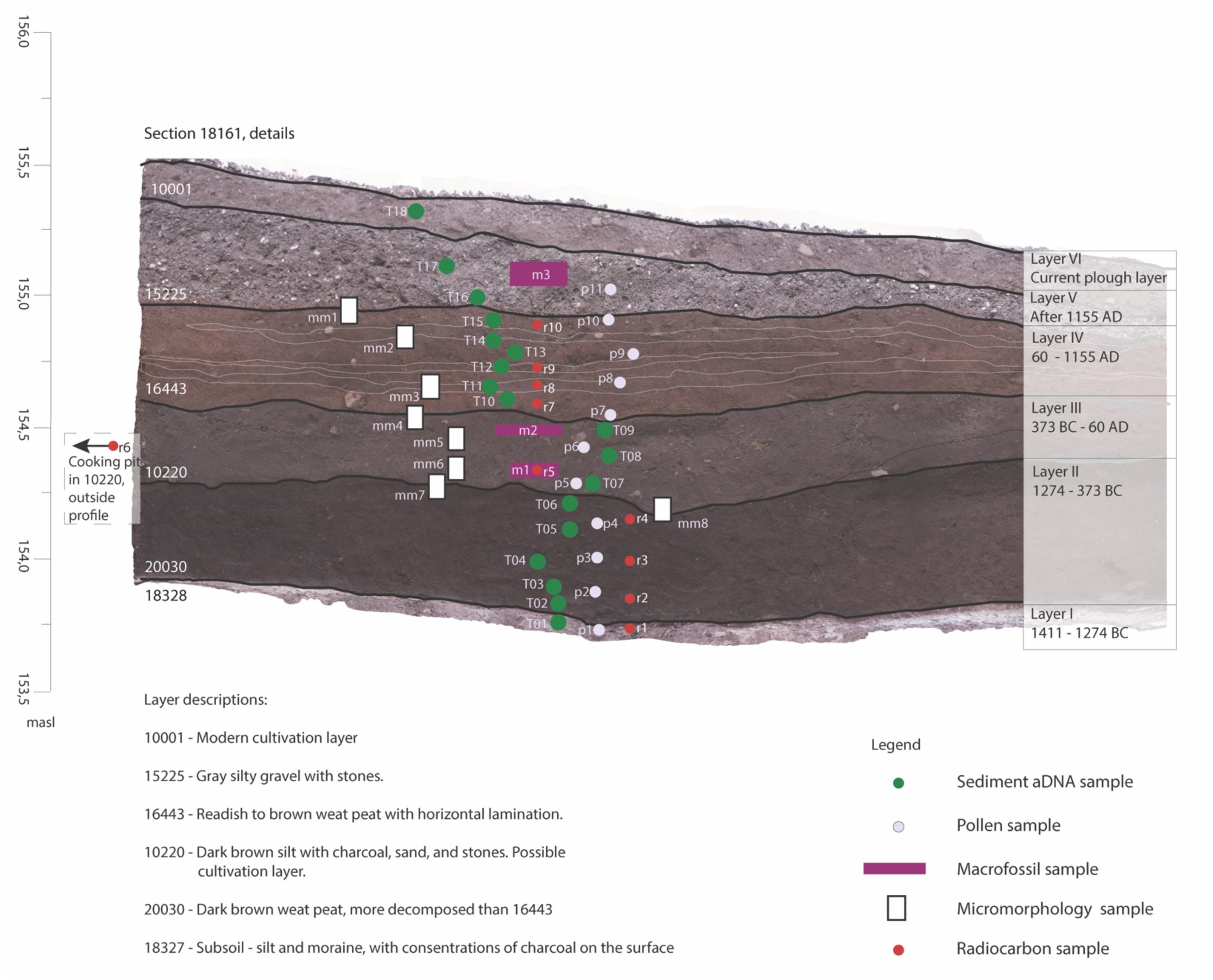
Profile showing sampling locations through Layers I-VI for sedaDNA (T), radiocarbon dating (r), micromorphology (mm), macrofossil (m) and pollen (p) analysis.

Initial on-site analysis of stratigraphy and dating indicated that the activity began in the Late Bronze Age, represented by peat-containing bog layers. There are indications of grazing, and several small stone cairns, one of which contained a now decayed organic container or vessel, may be traces of sacrifice in wet ground. The next phase dates to the Early Iron Age and represents an alternation between cultivation and other activities. A thick cultivation layer with cooking pits and a number of postholes at different levels within the cultivation layer, belong to this activity. The last phase dates to the Late Iron Age and the Middle Ages, and is represented by a new bog layer and a cultivation layer sitting on top of it. In connection with the bog layer, several wicker fences, cooking pits, a possible well, and parts of a boat were found. These findings raised the questions: 1) What processes have formed the layers? 2) Can the use of sedaDNA and complimentary soil analyses shed light on landscape use and cultivation through time?

For the purpose of the study, one profile section was chosen for all sampling (see Figure 2). Eighteen sedaDNA samples were retrieved throughout the six distinct layers. Eleven pollen samples were taken in close proximity (Figure 2). Three macrofossil samples were retrieved from the agricultural layers. Eight micromorphology samples were collected from strategic sections of the profile, together with nine samples for radiocarbon dating (Figure 2).

### 2.2. Radiocarbon dating

For radiocarbon dating, charcoal and plant remains were collected in different depths of the Layers I-IV aiming to construct an age-depth relation for the profile. Charred wooden remains underwent wood anatomy analysis to identify short-lived tree species so that the old-wood effect is minimised. The selected samples underwent an acid-alkali-acid (AAA) treatment to remove soil remains that come from more mobile materials in the soil. The cleaned samples were combusted and graphitized in preparation for measurement by accelerator mass spectrometry (AMS)^16^. They were measured at the Trondheim 1MV accelerator^17^. The results were calibrated using OxCal 4.4.4^18^ against the IntCal20 calibration curve^19^.

### 2.3. Soil micromorphology

To assess the site and soil formation processes, detailed geoarchaeological descriptions of two sections were undertaken, including documenting layer textures, interfaces, and composition, with reference to the wider topography. This was further aided by micromorphology, with eight samples assessed (Figure 2).

The samples for micromorphology were processed according to standard practice^20^, examined using plane polarised light (PPL), crossed polarised light (XPL) and oblique incident light (OIL) at x1 to x400 magnifications, and described using widely practiced methods^21–23^.

### 2.4. Pollen and macrofossil analysis

Samples for pollen analysis were collected from Layers I to V (Figure 2). In all, eleven samples were analysed. The samples were prepared following the procedures of Fægri & Iversen^24^. Briefly, humic acids were removed using 10% potassium hydroxide (KOH), minerogenic particles were removed with warm hydrogen fluoride (HF), and cellulose was removed via acetolysis. All samples were spiked with 3 tablets of *Lycopodium* spores and stained with basic fuchsin.

The results from the pollen analysis is given in a percentage pollen diagram constructed using Tilia 2.6.1^25^ (Supplementary Figure 2). The percentages are calculated from the total sum of pollen (ΣP), which is the sum of terrestrial pollen taxa and unidentified pollen. Spores, NPPs (non-pollen palynomorphs), and charcoal dust are kept outside of the total sum, and their percentages are calculated from the ΣP plus the count of each spore type/charcoal. The pollen taxa are sorted alphabetically in the following groups: trees, shrubs, dwarf shrubs, herbs, and unidentified.

Three samples were collected for macrofossil analysis, two from layer III and one from layer V (Figure 2). The samples were floated over sieves with mesh size 4, 1, 0.5, and 0.25 mm using tap water and analysed in wet state. To aid with the identifications the reference seed collection at the University of Bergen and Cappers et al.^26^ were used. The macrofossil results are given in a percentage diagram where the charred and the uncharred macrofossils are shown separately, with unique sums and percentages (Supplementary Figure 3).

### 2.5. Sedimentary ancient DNA methods

#### 2.5.1. Profile sampling

To reduce the risk of contamination of the sedaDNA samples, all sampling was performed using sterile personal protective layers, including hairnets, two pairs of nitrile gloves, sterile sleeves and facemasks. The outer layer of gloves was changed between handling each sample. Due to the pebbly nature of the sediments, coring was not possible. Rather, sampling was done using sterile 50 ml Falcon tubes that we inserted horizontally into the cleaned profile. The soil was crumbly, so to avoid soil from modern layers crumbling on the lower layer collection tubes, we staggered the points of collection (Figure 2). Additionally, large rocks distributed throughout the soil limited where the 50ml Falcon tubes could be placed. Sample T01 was a control sample taken from Layer I, which represented a layer deemed “sterile” by the archaeologists (meaning no human occupation was detected). For the bottom three cultural layers (Layers II, III and IV), samples were collected at approximately 5-cm intervals until sample T15. Samples T16 and T17 from Layer V were collected approximately 10 cm apart. Sample T18 represents the contemporary layer and is used as a modern control.

#### 2.5.2. SedaDNA extraction

DNA extractions and library constructions were performed in the positively pressurized paleo-genomic laboratory at the NTNU University Museum, which is specifically dedicated to aDNA analyses. Established criteria and rules for aDNA manipulation^27^ were followed during all the steps. All specimen tubes containing sediments were wiped down with 5% bleach then sterile water before being brought into the paleo-genomic laboratory. Tubes were opened only one at a time, and subsampled for aDNA extraction (Supplementary Table 2). Subsampling was performed by using a sterile pipette tip to scrape sediment from the middle of the tube, avoiding the sides and top layer.

Approximately 250 mg of sediment from each sample, along with two extraction blanks, were processed using a custom DNA extraction protocol based on the Qiagen DNeasy PowerSoil Pro Kit protocol. Specifically, 800 µl of Solution CD1 was added to the soil in PowerBead Pro Tubes (Qiagen) and the mixture was subsequently homogenized using the Qiagen LT TissueLyser at 25 Hz for 5 min. Next, 20 µl 1 M DTT and 20 µl proteinase K (20 mg/ml) were added to the homogenized sample, then mixed on the TissueLyser for an additional 5 mins at 25 Hz. The tubes were then incubated overnight at 37℃ on a shaking heat-block. The remaining steps were as per the manufacturer’s protocol until the addition of Solution C6. For this step, 45 µl of Solution C6 was added to the silica filter and incubated at 37℃ for 5 mins. The tube was then centrifuged at 15,000 rcf for 1 min. The flow-through was then re-incubated and centrifuged at the specified settings to increase the yield of extracted DNA. The DNA was sored at –20°C until downstream processing.

Given that 250 mg soil is a relatively small amount, and other methods reported in previous studies have used up to 5 g, we extracted two additional 250 mg sub-samples from four sample tubes, T01, T08, T13 and T16, as well as another extraction blank (QB1). These would be used to determine how robust the results from small portions of a larger sediment sample are. The two extraction blanks yielded no detectable DNA using the Invitrogen Qiagen HS 1x dsDNA kit.

#### 2.5.3. Library preparation and sequencing

Double-stranded sequencing libraries were prepared using the BEST method^28^. After adapter ligation, each sample was quantified using qPCR to determine the optimal number of cycles for the indexing PCR using AmpliTaq Gold polymerase (details of cycle numbers can be found in Supplementary Table 2). Libraries were then quality-controlled on the Agilent TapeStation 4100 using HS D1000 reagents to ensure correct molarity and fragment length for sequencing. Libraries were sent to the commercial service Novogene Europe (Germany) and sequenced on the Illumina NovaSeq X platform with 150-bp paired-end reads.

#### 2.5.4. Bioinformatic analysis

Sequencing reads were classified using an alignment-based holistic approach that uses sequences from multiple organisms with no prior assumption as previously described^29^. Adapters and low-quality bases were removed using cutadapt v4.9^30^ and AdapterRemoval v2.3.3^31^. Paired-end reads with a minimum overlap of 11 bp were collapsed, and then filtered for low-complexity reads and PCR duplicates using sga v0.10.15^32^. Filtered reads were competitively aligned to a selection of reference databases (NCBI RefSeq release 224, NCBI nucleotide collection downloaded on 10th November 2024, and Arctic-boreal plant and animal genomes), as described for the Holi pipeline^1,33^, using Bowtie2 v2.5.1^34^. In addition, the Genome Taxonomy Database (GTDB) was included in the reference databases^35^. A preliminary lowest common ancestor (LCA) was obtained for all reads using ngsLCA, and reads classified at levels higher than family were discarded from following analyses.

Alignments were merged, sorted and compressed before taxonomic classification. Multi-mapping reads were resolved and assigned to a unique reference sequence using bam-filter v2.0 (https://github.com/genomewalker/bam-filter). References were filtered using several metrics such as breadth of coverage (ratio between observed and expected > 0.8), normalized entropy (> 0.75) and Gini coefficient (< 0.6). The resulting alignment files were taxonomically classified to the LCA and damage rates for each taxa were estimated using metaDMG v0.4^36^. Read counts at the genus level derived from the alignments to the GTDB database^35^ were used to evaluate different environmental source proportions using SourceTracker2^37,38^, either using all counts or only those for authentic ancient microbes (minimum damage coefficient A > 0.02, and estimated significance Z > 2). Source datasets were derived from those used in Zampirolo et al.^8^.

## 3. RESULTS

### 3.1. Radiocarbon dating

A figure showing the calibrated dates from the profile can be found in Supplementary Figure 1. The on-site characterization of the layers indicated colluvial mass movement downslope, meaning that there was a degree of admixture, which varied by layer. There were also some mismatches in the dating sequence – the dating results from the two peat layers largely coincide, probably suggesting that the dating was done on material that has been washed into the deposits. The pollen content, presented below, in the two peat layers were so different that it seems unlikely that they are contemporaneous^39^. This is also supported by the geoarchaeological and soil morphology analyses, which demonstrates different formation processes^40^. These dates are therefore excluded from the discussion of our results, but included in the supplementary material (Supplementary Figure 1). The dates that were deemed most reliable are presented in the profile Figure 1.

### 3.2. Geoarchaeological analysis

A full report of the soil micromorphology analysis can be found in Supplementary Table 1. In summary, the sediment build-up was found to be a combination of anthropogenic and natural soils. On-site stratigraphic analyses revealed for the earliest layers an early peat formation after a shallow water body had been filled in with sediments, creating a highly organic, wetland soil. Later, the peat formation became more influenced by human activities in the area, such as land clearance and cultivation, which caused increasing colluvial sedimentation. The human influence, and the colluvium, are not steady; the speed of accumulation changes over time, suggesting a degree of temporal variation in human activity (Supplementary Table 1).

In detail, Layer I was deposited under shallow water, in moderate energy conditions, with seasonal/water level-based fluctuations in deposits. No root channels were visible, suggesting fine plant growth and/or that the organic matter came primarily from inputs into the water. Layer II formed as a shallow water body accumulated increasing organic matter and gradually moved from fenland to a transitional bog. The majority of the layer probably represents groundwater-fed, and thus fairly nutrient-rich, fenland, that accumulated slowly on the edges of a natural depression for which the silt beneath impeded drainage. Layer III represents a mix of peat formation due to the high water levels and topographic position, mixed with colluvial inputs from upslope from manured fields, and is distinct from the layers above and below. The higher mineral content in this context, compared to the contexts above and below, suggest that there was perhaps a shift in land use upslope which significantly affected soil formation and thus vegetation. In Layer IV, the peat formation looks to be primarily rainwater-fed but seems to have been subject to temporal change through inputs of colluvium and possibly changes in water levels. These changes are probably caused by human land use upslope, which affected the peat formation by increasing mineral inputs and the type of organic inputs. The upper half of the context shows that colluvial inputs were increasing in intensity and frequency, meaning the accumulation rate increased. There are also signs of a change in vegetation, from finer plant materials such as grasses and mosses at the base, to coarser, more woody plants at the top. Layer V is colluvial, with high initial accumulation rate, potentially slowing gradually over time. There is a very small transition to the peat below in terms of soil matrix mixing, suggesting that this is either an erosional boundary, or more likely that it represents a rapid change in soil formation caused by land use change. Lastly, Layer VI represents the disturbed, modern layer.

### 3.3. Pollen and macrofossil analysis

The pollen diagram (Supplementary Figure 2) shows a marked change in the pollen content between the different layers of the section. In Layer I, pine (*Pinus*) and birch (*Betula*) dominate the assemblage, and there are few herb taxa present. Presence of dropwort (*Filipendula*) and bog moss (*Sphagnum*) indicates wet soils, and there is less than 5% charcoal dust. In Layer II, the lower peat layer, the amount of tree pollen is high, dominated by birch (*Betula*) and alder (*Alnus*). Other tree taxa are present, but to a lesser extent, and only twelve herb taxa in the samples. There are low values for the dung indicating fungus *Cercophora* (HdV-112)^41^, and low occurrences of taxa indicating grazing. No charcoal dust has been recorded in the samples. Layer III, assumed to be an agricultural layer, displays a marked reduction in tree pollen. Total amount of herb taxa increases to 29 and the herb assemblage is dominated by Poaceae. Pollen from Poaceae indicates meadows or grassland areas. In addition, numerous arable weed taxa alongside cereal pollen of *Hordeum* and *Triticum* indicate the presence of arable land and cereal cultivation in the area. There are several taxa of dung indicating fungal spores in the pollen samples (*Sordaria, Cercophora, Sporormiella* and *Podospora*) indicating possible livestock grazing or manuring of the fields. The amount of charcoal dust reaches 70-80%. The macrofossil samples from this layer contain charred fragments of *Hordeum* kernels in addition to charred seeds of meadow plants. Several plants associated with water/wet soils are present in uncharred state (*i.e. Callitriche, Rorippa, Montia etc.*). There are also presences of both arable weeds and taxa indicative of grazed meadows present. In the topmost peat layer (Layer IV), the amount of tree pollen decreases further. The total amount of herb taxa increases to 39 in this layer, and is dominated by Poaceae and Cyperaceae. Meadow and grassland taxa are present. Cereal pollen of *Hordeum*-type is present in the bottom sample, and *Avena, Triticum* and *Secale-*type are present in the topmost sample. Arable weed taxa are also present. Dung indicating fungal spores of *Sordaria, Cercophora, Sporormiella* and *Podospora* are still present. Spores of the fungus *Clasterosporium caricinum* are also present, indicating that *Carex* taxa are represented within the Cyperaceae curve. The amount of charcoal dust varies between 20-60%. The topmost sample in the sequence from Layer V has low tree pollen, while herb taxa dominate the assemblage with 26 herb taxa identified, and meadow and grassland taxa are still present. Of these, Poaceae dominates and there are increases in the arable weed taxa. Cereal pollen of *Avena* and *Hordeum*-type are present with maximum values of the sequence. There are increases in the dung indicating fungal spores of *Sordaria* and *Sporormiella.* Charcoal dust increases to over 80%. The macrofossil sample from layer V shows a dominance of seed of the Caryophyllaceae, *Spergula* as well as a high proportion of unidentified seeds. Generally, the seeds in this layer are from arable weeds and meadow plants.

### 3.4. SedaDNA metagenomic analysis

#### 3.4.1. Sequencing data summary

Metagenomic DNA was obtained and processed into 22 samples and 2 negative control libraries, resulting in a total of 1,858,602,644 150-bp read pairs. For all samples, the majority of the data was composed of adapters and low quality bases, low complexity reads and duplicates (Supplementary Figure 4). 12.57 Gbp of sequencing data was retained from the total as usable for taxonomic classification. A complete summary of read statistics in each step of the filtering process can be found in Supplementary Table S2.

As expected, only a small fraction of sample reads were successfully classified following the analyses. The percentage of reads entering the analysis that were successfully classified to any organism ranged from 0.055% (1,524 reads) for sample T01.2 to 1.82% (177,368 reads) for sample T13.2. Although some reads from extraction blanks were initially classified, none of these passed the alignment filtering. Given the complexity of species-level identification in environmental DNA samples, we predominantly present genus-level classifications. Most classifications were bacterial, with a total of 1,009,778 reads assorted into 962 different genera. Another 406,928 reads were assigned to 55 genera within Archaea. For eukaryotes (any organism that possesses a clearly defined nucleus), a total of 98,893 reads were classified to 113 different genera, although, as it will be discussed further below, roughly half of the reads (48,820) were likely-modern *Hordeum* fragmented reads from the topmost layer (T18). Plants were the most diverse phylum with 91 different plant genera to which 96,198 reads were classified, followed by animals with 443 reads assorted into 10 animal genera. Lastly, sample T05 resulted in no reads classified as a eukaryota genus, despite having a similar number of total classified reads as the other samples. The lack of identifications for this sample may be due to it being not representative enough of the layer, or due to specific soil properties not optimal for enough aDNA preservation as to enable eukaryota identifications.

To further understand the influence of processing a relatively small amount, 250 mg, from a 50 ml falcon tube in the Eukaryota detection and profiles, we included two replicates for samples T01, T08, T13 and T16. For plants, replicate profiles remained similar for samples T01, T13 and T16; while for T08 the replicates differed in taxa composition (Figure 3A). For animals, neither replicates for samples T01 and T13 yielded animal reads, and replicates for samples T08 and T16 were different both in classification diversity and read abundance (Figure 3B). The similarity between plant replicates compared to the distinct animal profiles was expected due to the nature of their DNA deposition.

**Figure 3.**
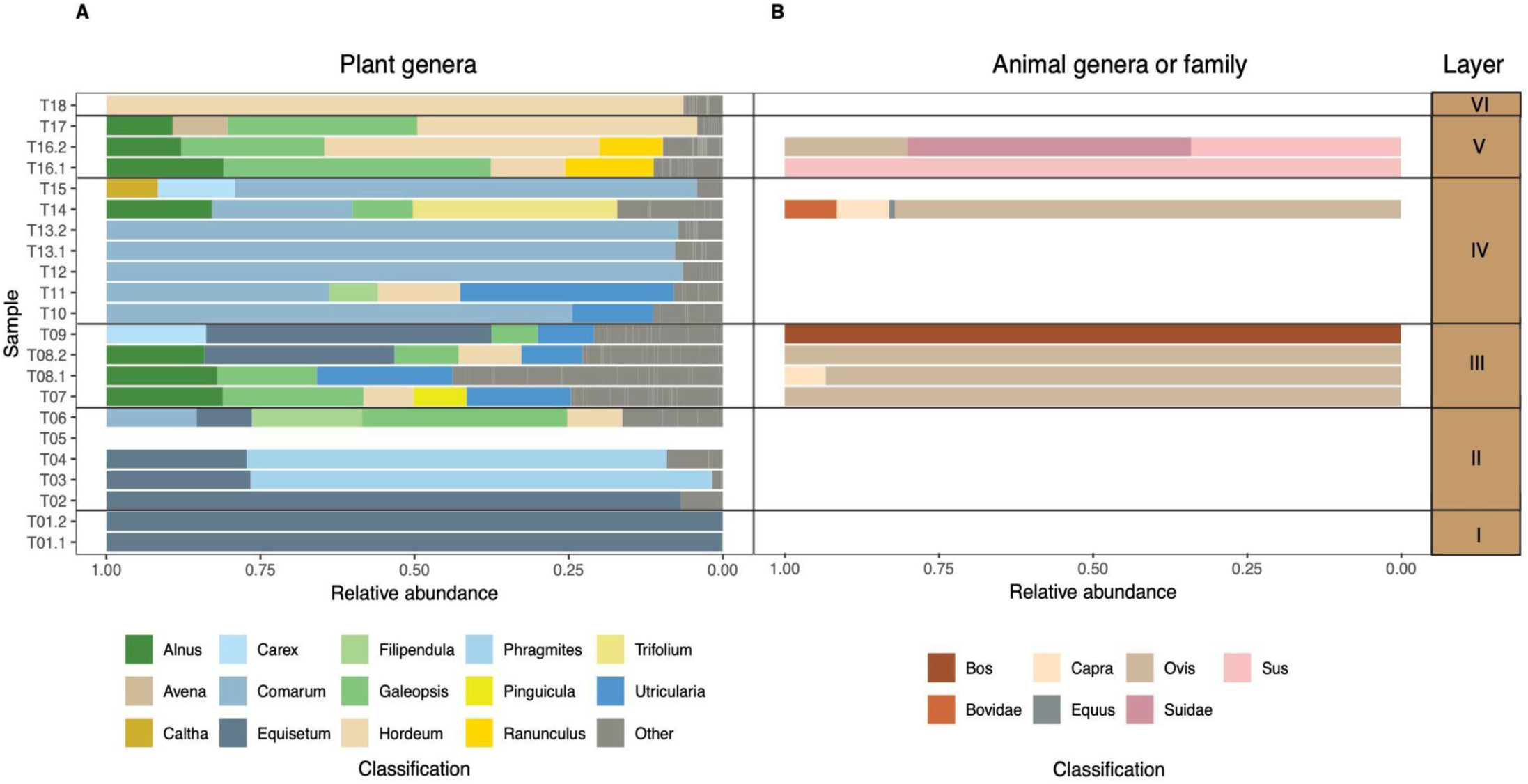
Relative abundance plots in each sample in Layers I-VI for A) Plants, where genera with an abundance under 5% were classed as “Other”; and B) Animals at either the genus or family levels.

#### 3.4.2. Plant detection

Plants were the most diverse and abundant Eukaryota classifications, including 216 different species, 92 genera, 32 families and 20 orders detected (see Supplementary Table S3). As indicated above, we discuss classifications at the genus level as species-level identifications may be greatly confounded by taxonomic ambiguity and/or database underrepresentation^42^. Genus-level calls were manually checked to determine whether they were likely to be present in Norway. Seven of the 92 genera were determined to be genera not found in Norway, and as such were called at the family level in Figure 3A. These included *Aegilops*, *Agropyron*, *Andropogon*, *Cryptomeria*, *Dasiphora*, *Helictochloa*, and *Zea* Of note, *Zea* reads were detected in moderate numbers, representing 2% of the plant reads overall. *Zea* is not a crop that has been grown in pre-historic Norway hence we omitted it from the Figure 3A on the assumption it could represent either a reference contamination or a closely related grass species that is not represented in the database.

In Layer I, *Equisetum* dominates in both replicates, with small amounts of *Avena*, *Galeopsis* and *Carex* in one of the replicates. The presence of *Avena* in one of the replicates was not expected, and likely indicates that this signal is contamination between samples (lab contamination) or naturally occurring soil mixture (transition layer contamination). *Carex* next occurs in sample T02 (bottom of the next layer), *Galeopsis* next appears in the top of Layer II (sample T06), while *Avena* does not appear until Layer IV (sample T07). As such, we deem these genera to be unreliably present in Layer I.

In Layer II, sample T02 is still dominated by *Equisetum* with low levels of *Carex*, but by sample T03 the profile shifts slightly to be dominated by *Phragmites* with low levels of *Hordeum*. This is the first occurrence of a crop genera in the profile. *Hordeum* then drops out until sample T06, the top most sample of this layer. Sample T05 had no reads passing filter. Overall, this layer has few genera detected (<5) in all samples except the topmost sample, T06, of this layer indicating an increase in biodiversity by Layer III. In sample T06, we see a change in the dominant genera from *Equisetum* and *Phragmites*, wetland-associated plants, to *Galeopsis* and *Filipendula*, both plants associated with drier soils.

In Layer III, the number of plant genera increased substantially indicating a shift in land use. The dominant genera in this layer include *Galeopsis*, *Equisetum*, *Utricularia*, *Pinguicula*, *Carex*, *Filipendula*, as well as *Hordeum* (barley) and low levels of *Avena* (oat) and *Humulus* (hops). These genera include a mix of wetland/marsh (e.g. *Equisetum*, *Utricularia* and *Carex*) plants, as well as more genera indicating crop agriculture (e.g. *Galeopsis*, *Filipendula*, *Hordeum*, *Avena* and *Humulus*) and, surprisingly, *Alnus*, a tree which was indicated as decreasing in prevalence based on the pollen counts.

In Layer IV, samples T10 and T11 the plant genera become dominated by *Comarum*, *Equisetum*, *Utricularia*, and *Carex*. Sample T10 contains very low amounts of *Avena*, while T11 is the only sample in this layer with relatively high abundance of *Hordeum* (Figure 3A). which are all marsh-associated plants, and the number of reads associated with crops decreases. Samples T12 and both T13 replicates show a marked reduction in cultivated taxa, with *Comarum* dominating. The top portion of this layer, including samples T14 and T15, show an increase in meadow-associated taxa such as *Trifolium*, *Filipendula* and *Caltha*. Interestingly *Alnus* appears in high abundance in this layer, coinciding with the re-emergence of domesticated animal species.

By Layer V, *Hordeum* dominates the plant taxa, along with meadow-associated taxa such as *Ranunculus* and *Filipendula*. *Alnus* remains abundant throughout this layer.

Layer VI, representing the modern topsoil, is dominated by *Hordeum*, with some *Avena* and *Triticum* also detected. *Galeopsis*, *Ranunculus*, *Alopecurus*, and other meadow taxa are detected in relatively lower abundances.

#### 3.4.3. Animal detection

In this study, only a small portion of the analyzed reads were confidently classified as animal genera. These included seven in the class Mammalia (*Equus*, *Bos*, *Ovis*, *Budorcas*, *Sus*, *Phacochoerus* and *Capra*), two Insecta (*Condylostylus* and *Sitodiplosis*) and one Gastropoda (*Pomacea*). From these, *Phacochoerus* and *Budorcas* were deemed unlikely to be true calls and were depicted at the family level Suidae and Bovidae, respectively, in Figure 3B. It is likely that these reads correspond to regions shared between closely related genera and thus were spuriously classified during the alignment reassignment and filtering step, as has been done in similar studies^8^. Only mammals were plotted, being the most relevant taxa to this study.

While no animals were detected in Layers I and II, all samples taken from Layer III detected mammals. The bottom two samples, T07 and both T08 replicates, detected *Ovis*, in addition to T08.1 detecting low levels of *Capra*. The topmost sample, T09, contained a relatively high number of *Bos* sequences. In Layer IV, only one sample, T14, detected mammals. This was at the top portion of the layer. Interestingly, this sample contained the most mammal diversity with *Ovis*, Bovidae, *Capra* and *Equus* detected. Layer V, in the lower sample, detected *Sus* in T16.1 and *Sus*, Suidae and *Ovis* in the replicated T16.2 sample.

#### 3.4.4. SourceTracker microbial analysis

Output from SourceTracker can be found in Supplementary Figure 5A and 5B. The output text files from SourceTracker were manually edited to combine similar groups, this included combining: Melting permafrost and Permafrost, *Ovis* dung and *Ovis* rumen, *Bos* dung and *Bos* rumen, Freshwater sediment and Lake in Figures 4 and 5. Additionally, to gain greater insight into the patterns of the known source microbes, plots were made excluding the Unknown category (Figures 4B and 5B). Following the approach of Zampirolo et al.^8^, we plotted all microbes (Figure 4) and only microbes determined to be ancient (Figure 5), as both reveal relevant information about the soil composition through time.

**Figure 4.**
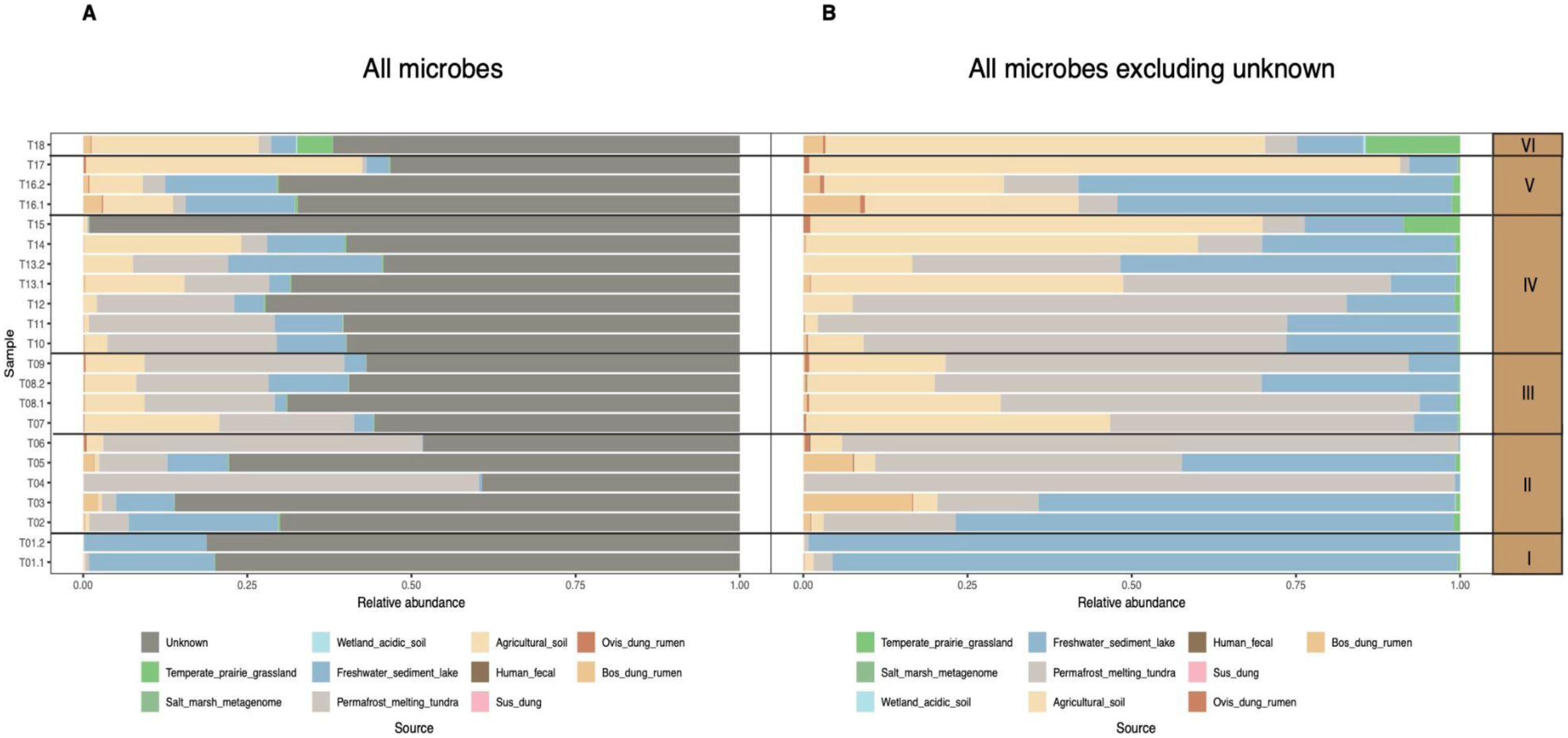
SourceTracker microbial characterization for A) All microbes; and B) All microbes excluding the Unknown category.

**Figure 5.**
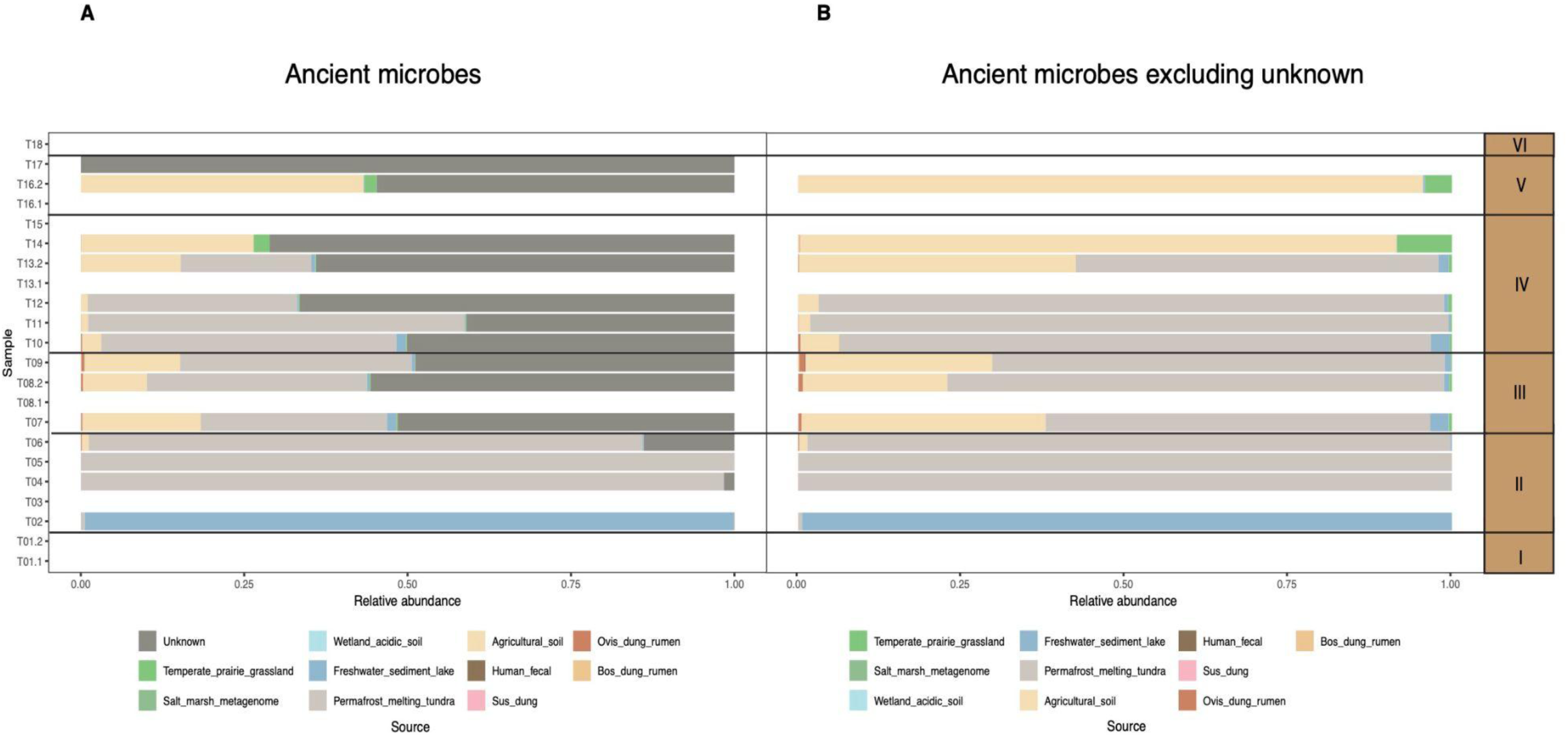
SourceTracker microbial characterization for A) All microbes deemed to be genuinely ancient microbes; and B) microbes deemed to be genuinely ancient excluding the Unknown category.

When considering all microbes (ancient and modern), the dominant sources other than Unknown (a likely reflection that the database we used contained few directly relevant microbes to Norwegian soil) are freshwater sediments and lakes, permafrost and melting tundra, and agricultural soil. It should be noted that Torgård, even 3500 years ago, was not considered tundra or permafrost. Unfortunately there are few perfectly suited reference databases for this location. As such, we interpret permafrost– and tundra-related microbes to signify simply cold conditions at our site. Interestingly, the ancient microbes contained very little freshwater sediment and lake associated microbes, with the exception of sample T02.

In terms of animal associated microbes, *Bos* dung and rumen associated microbes were detected in the all microbe dataset in Layers II and V most dominantly, and in lower levels in Layers III, IV and VI. *Ovis* dung and rumen associated microbes were detected in very low levels throughout the layers. Of the ancient microbes, low levels of *Ovis* dung and rumen associated microbes were detected in the top of Layer II and throughout Layer III. This was consistent with the appearance of domesticated animals in the layer. Interestingly, Layer V revealed the second most abundance of Bos dung and rumen associated microbes in the All microbes dataset, while not being detected in the ancient despite domesticated animals being detected in this layer. This could indicate that the microbes were degraded to a lesser extent in these upper layers.

## 4. DISCUSSION

The application of sedaDNA metagenomic analysis at Torgårdsletta reveals a distinct temporal progression in plant, animal, and microbial communities, enhancing our understanding of the interplay between human practices and environmental changes in central Norway. While this dataset is representative of a single site, it provides valuable insights into land use patterns from the late Bronze Age to Medieval times in Norway, specifically highlighting the transition from peat bogs to arable land for crop cultivation and livestock grazing. This study exemplifies how sedaDNA can complement traditional palaeobotanical and archaeological proxies, deepening our comprehension of anthropogenic impact on the evolving landscape of the region. In this section, we will explore the transitions over time from the lowest identified layer (Layer I) to the uppermost historical layer (Layer V), focusing on how these changes reflect shifts in land use.

### 4.1. Early Bronze Age

Layer I, the deepest layer in the Torgårdsletta profile, dates to 1411-1279 BC, corresponding to the early Bronze Age. This layer is characterized as a minerogenic layer with silts and stone (moraine), with minimal archaeological evidence indicative of human activity. The low levels of charcoal dust observed in the pollen analysis, along with a marked absence of plant diversity in both pollen and sedaDNA analyses, support this characterization. The sedaDNA results reveal limited plant diversity, aligning with the geoarchaeological assessment that suggest the layer formed under shallow water conditions. Equisetum was the dominant plant identified in Layer I, however, the observed low plant diversity may also reflect a lack of preserved ancient DNA in this lower layer. The DNA damage patterns derived from *Equisetum* indicate that the sequences obtained are genuinely ancient. However, the microbial DNA analysis resulted in no ancient microbes being identified, further supporting the hypothesis of poor DNA preservation in this layer. Further investigation of Bronze Age and older sediments from central Norway may provide insights into the preservation potential of ancient DNA in these contexts. Overall, the total microbial content in Layer I aligns with freshwater lake sediments rather than agricultural soil, reinforcing the interpretation that this site was not extensively cultivated during the early Bronze Age.

### 4.2. Late Bronze Age to Early Iron Age

Layer II, which has been dated approximately to 1274 – 373 BC, based on radiocarbon dates from the layers above and below, represents a gradual transition from woody, wet fenland to a mildly acidic bog with grasses and mosses, as evidenced by geoarchaeology and micromorphological analysis. Archaeological excavations in 1998 identified significant settlement activity on the western slope of Torgårdsletta from around 1000 BC to 100 BC, with an estimated 70 households reported^13^. Grønnesby noted a rapid increase in the number of cooking pits at the site between 700 BC and 100 BC, followed by a decline. These cooking pits, located in the transition zone between dry land and marshland, suggest rituals associated with food preparation in this ecological boundary^43^. Microbial sedaDNA data indicate a transition from primarily freshwater lake-associated microbes from Layer I to an increase in agricultural soil and cold soil-associated (referred to as permafrost/tundra based on the reference database used in Figures 4-5) microbes in Layer III, reflecting ecological changes across the site. The ancient microbial sources indicate the first appearance of agricultural microbes in the uppermost sample of this layer. Additionally, transitions in plant communities, as revealed through pollen and sedaDNA analyses, show a shift from shallow water conditions to groundwater-fed environments, with aquatic plants such as *Phragmites* dominating the lower portion and an increase in *Galeopsis* (specifically *G. tetrahit*), a plant indicative of disturbed soils, towards the top. The sedaDNA analysis also reveals an increase in plant diversity, commonly associated with periods of land clearing^44^.

Small quantities of *Hordeum* (barley) were detected within this layer. During the Late Bronze Age, *Hordeum* was among the dominant grains grown in Norway^45^, so its presence, though limited, may suggest that it was cultivated nearby. However, the DNA damage profile does not confirm the authenticity of the *Hordeum* as genuinely ancient, raising the possibility that this genus could represent contamination from higher layers. The absence of pollen or macrofossil evidence to support the presence of *Hordeum* casts further doubt on its significance in this layer. However, considering that cereal pollen grains have low dispersal capacity^46^, it is plausible that the sedaDNA is detecting *Hordeum* grown at some distance to the site.

No sedaDNA from animals were found in this layer, which is consistent with the lack of bone material detected prior to c. 50 BC in the Ørland excavations off the coast of Trondheim. Perhaps this indicates similar preservation conditions or even a limited amount of animal husbandry occurring in a fixed location during this period.

### 4.3. Pre-Roman Iron Age

Layer III is dated to the Pre-Roman Iron Age (approximately 373 BC to 60 AD). This period sees significant developments in farming practices across Scandinavia. During this period, agricultural practices developed as farms grew larger, establishing centers of regional power. The introduction of longhouses, such as the Bronze Age/Early Iron Age structure located approximately 50 meters northwest of this profile, facilitated the protection of livestock during harsh winters^47,48^. Mixed farming became the primary mode of food production in southern Norway during the Early Iron Age, encompassing crops like *Hordeum* (barley), *Triticum* (wheat), and, to a lesser extent, *Avena* (oat), although oat is not regarded as a substantial crop during this time^49,50^. In this region, commonly raised livestock included cattle, sheep, goats, pigs, and horses.

The evolving agricultural landscape is evident in both the sedaDNA and other proxies, with soil micromorphological analysis indicating an increase in mineral content in the upper layers and a marked shift in vegetation. Both pollen and sedaDNA analyses reveal a rise in herbaceous plants and grasses, including barley and wheat in the pollen sequence, along with an increase in plant diversity suggesting land clearance and cultivation. Interestingly, the sedaDNA analysis detected very low levels of *Humulus* (hops), which is significant given that hops, predominantly used for beer brewing, indicates an important aspect of cultural practices. Local hemp/hops retting in Scandinavia is suggested to have begun around AD 1-400^51^, and this is further supported by pollen analyses from the Eidsvatnet site, located approximately 90 km northeast of Trondheim, during the same period^52,53^. However, it is important to note that pollen analysis cannot easily differentiate between hemp and hops (*Cannabis*/*Humulus*) when preservation conditions are poor^54^. Thus, these sedaDNA sequences represent the earliest and first direct evidence of hops cultivation in this area. Given the low read count for this genus in the sedaDNA results, further sampling from additional profiles at the site would be necessary to validate this finding definitively. Notably, there are key differences between the sedaDNA and pollen data in Layer III, underscoring the complementary nature of these methodologies. While pollen analysis consistently identifies *Alnus* (alder) and *Betula* (birch) across all layers, there is a significant decrease in their pollen representation inLayer III. In contrast, sedaDNA analysis shows the presence of *Alnus* and *Betula* starting from Layer III, with *Alnus* being particularly dominant. This discrepancy is probably an indication of reduced forest cover, as the land was cleared for farming and grazing, and the sedaDNA may reflect the local use of *Alnus* for livestock fodder or wood-working, with fences and wooden wells detected from around 458 – 529 AD. Alnus charcoal were also detected in cooking pits, where some date to the same period.

Additionally, we observe a distinct increase in aquatic and graminoid plant genera in the sedaDNA for Layer III compared to earlier layers, showcasing greater diversity relative to the pollen findings (Supplementary Figure 4). While both proxies indicate that this layer was agricultural, they differ in their representation of overall plant diversity. The pollen analysis predominantly characterizes the landscape as open grassland or meadows. Similarly, the sedaDNA shows a transition to grasslands with higher taxonomic resolution of Poaceae detected than in the pollen data. Moreover, sedaDNA analysis suggests an increase in plants associated with wet and nutrient-rich soils, as well as increased DNA reads associated with tree taxa. These differences may arise from the limitations of pollen preservation or the lack of taxonomic resolution to identify graminoids into species, with the exception of cereals and Phragmites^55^.

SedaDNA analysis further reveals a transition from predominantly cold soil– and freshwater-associated microbes to a notable increase in abundance of agricultural soil-associated microbes.Pollen analysis identifies several fungal spores indicative of dung, andsedaDNA indicates the first presence of mammals, specifically sheep, goats, and cattle. The lower portion of this layer suggests the presence of sheep and goats, with cattle detected only in the upper portion. Although the low number of reads restricts definitive conclusions regarding prior livestock presence, the timing of their detection aligns with the land-use changes indicated by the other proxies.

Moreover, although detected at low levels, ancient sheep gut/dung-associated microbes were detected throughout this layer, reinforcing the notion that livestock first appeared at this site during the Early Iron Age. This finding is significant due to the scarcity of osteological remains at this site and others in central Norway from this period. The livestock genera detected correspond well with findings from a nearby Pre-Roman Iron Age site (400 BC – 0) in Ørland, where a preserved bone midden contained similar livestock species (sheep, goats, pigs, horses, and cattle), suggesting that our findings reflect broader trends in animal husbandry at farm sites throughout central Norway during this time^56^. The observed transition in land use and the introduction of livestock at this site coincide with changes in farming practices accompanying the advent of iron sickles around 200 BC^57^, marking a shift from forest pastures to animal stalling and more localized grazing. The sedaDNA evidence of livestock presence supports the hypothesis that these animals were initially forest-grazing, complicating the detection of their DNA unless samples were taken from areas that were previously forested.

### 4.4. Iron Age to Medieval Period

In Layer IV, between approximately 60 to 1155 AD, land use at the site underwent a notable transition from agricultural soils to peat horizons, as indicated by geoarchaeological analysis in Layer IV. This analysis reveals a change in vegetation throughout the layer, with grasses and mosses dominating the lower portion, while woodier plants become more prevalent toward the top. Of note, pollen and sedaDNA analyses identify the presence of barley, wheat, and oats in the lower section, but not in the upper portion of the layer, indicating that there was a decrease in agricultural crops grown in the vicinity during this time. Additionally, there are increases in *Carex* and *Sphagnum* as well as presence of *Potamogeton* in the pollen analysis, and *Comarum* (specifically *C. palustre*) dominating the sedaDNA plant reads in this layer.

This shift in vegetation is accompanied by shifts in the sedaDNA microbial signatures. In the lower portion of Layer IV, the sedaDNA analyses indicate the presence of plant species associated with wet peat soils, with microbial profiles showing an initial increase in freshwater lake-associated microbes in the lower section of the layer, alongside increasing cold soil-associated microbes. Conversely, in the upper portion, these signals decrease while microbes linked to agricultural soils increase, suggesting significant ecological changes as well as changes in land use, supported by a shift in plant taxa associated with meadows and land clearing.

Pollen analysis detected spores associated with dung, while sedaDNA revealed limited amounts of microbes linked to animal dung in this layer, suggesting a local decline in livestock activity at the site during this time. In contrast, evidence of livestock was only detected in the upper portion of the layer (sample T14), which also corresponds with an increase in microbes associated with agricultural soils. Here, the highest diversity of animals—including sheep, goats, horses, and unspecified Bovidae—was observed. Notably, the presence of horses in this upper section aligns with dates from a nearby Vendel-type elite equestrian warrior grave excavated at Torgårdsletta in 1933^15^, possibly highlighting the increased usage of horses at the site during this time.

Of ecological importance, the shift of vegetation and land use indicated by the various proxies in Layer IV, aligns with the climatic events of AD 536 and AD 539-540, which were caused by volcanic eruptions and resulted in a marked cooling of the climate during the mid-sixth century^58–60^. Earth System Model simulations for southern Norway indicate air temperatures dropped by up to 3.5°C during this period^61^, with the years between AD 536 and AD 545 recognized as the coldest decade within that time frame^59,62–64^. These climatic changes may have driven the observed alterations in vegetation^61^. The changing climate coincided with shifts in societal structure, raising questions about whether the observed changes in vegetation and soil types were a direct result of the climatic conditions or reflective of alterations in farming and land use practices, including a need for water for animal husbandry. This would align with geochemical and palynological analyses of sediments in bogs, lakes and archaeological sites, which indicate that communities adjusted to the cooling climate post 536 AD by taking up hardier crop species and by shifting their focus to animal husbandry^65,66^. More precise dating information would be needed to reliably determine the cause of the ecological changes detected within this layer.

### 4.5. Medieval Period

Layer V, dating from approximately 1155 AD to approximately 1850 AD, has been identified as an agricultural layer due to the increased prevalence of cereals—such as barley, wheat, and oats—evident in both pollen and sedaDNA analyses. The microbial analysis via sedaDNA indicates a significant shift from cold soil-associated microbes in Layer IV to an increase in freshwater lake-associated microbes in the lower portion of Layer V, culminating in the dominance of agricultural soil-associated microbes in the upper section. Additionally, geoarchaeological analysis suggests that the layer experienced rapid colluvial deposition, potentially attributed to upslope, as suggested by the microbial indicators of freshwater lake sediments.

In terms of livestock evidence, pig and sheep were detected in replicates from the lower portion of this layer, whereas no animals were identified in the upper section. While microbes associated with pig feces were not found at meaningful levels in the bottom portion, microbes associated with *Bos* and *Ovis* dung were present.

## 5. CONCLUSIONS

The overall data leads to the interpretation that use and organization of fields during the Early Iron Age were likely more shifting compared to those in the Late Iron Age and Middle Ages. It appears that the establishment of the permanent agricultural fields we recognize today emerged following the transition to the Late Iron Age. This shift toward land ownership as a fundamental legal principle in society is essential for structuring the landscape effectively. The flexibility observed in settlement and farming practices, particularly during the Late Bronze Age and Pre-Roman Iron Age, was incompatible with established principles of land ownership. In locales where cultural layers have been preserved in stratigraphic sequences, the unstable patterns of settlement and agriculture can also be discerned later in the Early Iron Age.

In conclusion, our study reveals how metagenomic analysis of sedaDNA provides additional insights into the intertwined relationships between animal exploitation, landscape utilisation, climatic conditions and evolving societal needs. This genetic information complements the pollen, micromorphological and macrofossil records with a more nuanced representation of various genera, (plants but also animals) facilitating assessment of the complex interactions between the environment and human impact and agricultural practice. Furthermore, this study highlights the promising use of sedaDNA at an open air site that has experienced millennia of shifty climatic conditions, environmental development and human cultivation and activities, suggesting that DNA could be preserved in detectable amounts in many more archaeological sites than originally thought. Wider application of such methods could provide many more reference points like Torgårdsletta and help to further unravel the coevolution of farming practices in relation to climatic changes in Scandinavia and northern Europe. Sediments within archaeological contexts should be regarded as essential genetic archives for comprehending the complexities of agricultural developments in prehistory.

